# Targeting tumor-intrinsic metabolic node sensitizes pancreatic cancer to anti-PD1 therapy

**DOI:** 10.1101/519462

**Authors:** Nikita S Sharma, Vineet K Gupta, Vanessa T Garrido, Roey Hadad, Brittany C Durden, Kousik Kesh, Bhuwan Giri, Anthony Ferrantella, Vikas Dudeja, Ashok Saluja, Sulagna Banerjee

## Abstract

Pancreatic ductal adenocarcinoma (PDAC) is considered to be a highly immunosuppressive and heterogenous neoplasm. Despite improved knowledge regarding the genetic background of the tumor and better understanding of the tumor microenvironment, immune checkpoint inhibitor therapy (targeting CTLA-4, PD-1, PD-L1) has not been very successful against PDAC.

The robust desmoplastic stroma, along with an extensive extracellular matrix (ECM) that is rich in hyaluronan, plays an integral role in this immune evasion. Hexosamine biosynthesis pathway (HBP), a shunt pathway of glycolysis, is a metabolic node in cancer cells that can promote survival pathways on one hand and influence the hyaluronan synthesis in the ECM on the other. The rate-limiting enzyme of the pathway, glutamine-fructose amidotransferase (GFAT1), uses glutamine and fructose 6-phosphate to eventually synthesize UDP-GlcNAc.

In the current manuscript, we target this glutamine-utilizing enzyme by a small molecule glutamine analog (6-diazo-5-oxo-L-norleucine or DON). Our results show that DON decreases the self-renewal potential and metastatic ability of tumor cell. Further, treatment with DON results in a decrease in hyaluronan and collagen in the tumor microenvironment, leading to an extensive remodeling of the ECM. this in turn, increases CD8+ cytotoxic T-cells infiltration, and makes the tumors tumors more amenable and sensitive to anti-PD1 therapy.

## Introduction

Pancreatic cancer is a devastating disease with very poor outcomes. In spite of concerted efforts to develop effective therapeutic strategies, the five-year survival rate of the patients diagnosed with this disease is a mere 7% (1, 2). High rates of tumor relapse, therapeutic resistance and metastatic spread of the tumor contribute to these dismal statistics (3, 4). Compounding this, the pancreatic tumor is largely immune evasive. This makes these tumors resistant to immune checkpoint inhibitor therapies (with anti-PD1, anti-PDL1 and anti-CTLA4 therapy) that have shown remarkable benefit in other tumors (5, 6). Resistance to these immune therapy is predominantly due to the presence of an extensive fibro-inflammatory and desmoplastic stroma with a rich extracellular matrix (ECM). Together, these components of the tumor provide an immune suppressive microenvironment that prevents the infiltration of anti-tumor immune cells like CD8+ T-cells (7, 8). Targeting the stromal component (like activated stellate cells, cancer-associated fibroblasts or the hyaluronan-rich extracellular matrix) has shown sensitization of pancreatic tumors to standard chemotherapy over recent years (9–12). However, it is acknowledged that an anti-stromal therapy by itself is likely to result in a more aggressive tumor by relieving pressure on blood vessels and promoting metastasis (13). Thus, a treatment strategy that can deplete the stroma while eradicating the tumor cells in order to prevent metastatic spread is ideal for this devastating disease.

While the role of stellate cells in secretion of the robust ECM is well-known (7), how the ECM may contribute to the immune suppressive microenvironment within pancreatic tumor remains an enigma. Studies have shown that infiltration of CD8+ T-cells is associated with better patient outcomes (14, 15). Similarly, depletion of CD4+ T-cells in pancreatic cancer mouse models has been shown to result in decreased infiltration of several tumor-promoting myeloid cell population like macrophages and immature myeloid cells or myeloid-derived suppressor cells (MDSCs), indicating that the immune cell behavior within the tumor microenvironment is a complex interaction between different cell types (16) (17). In patients with pancreatic cancer, an immune response is typically mediated by CD8+ T-cell infiltration. However, a large number of these cells become trapped in the peritumoral stroma and do not reach the tumor to elicit an anti-tumor response (18). Migration of infiltrating T-cells is typically guided by stromal chemokines and ECM proteins (19, 20). Additionally, T-cells use the ECM fibers to migrate to tumor cells whether by ameboid-like contact guidance or using integrin-based adhesion (20, 21). Thus, remodeling of ECM is likely to have a profound effect on T-cell infiltration and function in pancreatic cancer (22).

The ECM of pancreatic tumors is primarily composed of glycosaminoglycans like hyaluronan (HA). HA is a polymer of glucuronic acid (GlcUA) and N-acetylglucosamine (GlcNAc). Synthesis of HA is orchestrated by HA synthases (HAS 1-3). These enzymes require UDP-GlcNAc as one of the primary substrates for synthesis of HA (23, 24). In a cell, UDP-GlcNAc is synthesized via the hexosamine biosynthesis pathway (HBP), a shunt pathway of glycolysis that utilizes glutamine and glucose to make this nucleic acid sugar (25, 26). Since HBP is also a nutrient sensing pathway, the synthesis of HA in these tumors is tightly coupled with the metabolic status of the cells. The rate-limiting enzyme of this pathway is glucosamine-fructose amidotransferase (GFAT1/2), which is responsible for controlling the flux of metabolites through this pathway. The HBP is highly utilized in a number of cancers, and its inhibition by either targeting GFAT1/2 or by preventing utilization of glutamine (using a glutamine analog 6-diazo-5-oxo-L-norleucine or DON) results in tumor regression (27–29).

UDP-GlcNAc is a major metabolite that is used in cellular glycosylation reactions. Decreased UDP-GlcNAc production can induce ER stress in the cells by inhibiting these reactions (30). Further, since UDP-GlcNAc is used by O-GlcNAc transferase (OGT) to glycosylate and regulate a large number of oncogenic proteins (like Myc), inhibition of UDP-GlcNAc production will affect the survival of the cancer cells (31). Consistent with this, previous studies from our laboratory have shown that HBP-fueled UDP-GlcNAc synthesis can be used by OGT to drive tumor growth and self-renewal (32, 33). Thus, HBP forms an integral metabolic node, inhibition of which will affect the tumor and its microenvironment alike.

While the role of HBP in cancer cell survival is being studied by several groups, whether HBP can be targeted genetically or pharmacologically to remodel the ECM, and thus make the immune evasive pancreatic tumor to immune therapy, has not been studied before. Since the hexosamine biosynthesis pathway is dependent on glucose uptake and glutamine equally, inhibition of glutamine utilization by using glutamine analogs like azaserine and 6-diazo-5-oxo-L-norleucine (DON) has been used over the years to target this pathway. Even though studies have shown that DON may have pleiotropic effects (34), it was used clinically as an anti-tumor agent (35). DON was successfully used in five phase 2 clinical trials (34), leading to disease stabilization. However, in spite of showing promise, DON was abandoned clinically as an anti-tumor agent. With the recent knowledge that tumor microenvironment is an integral component that drives progression of the tumors, there has been a renewed interest in reviving DON as an anti-tumor agent. Since recent evidence showed that a broad-spectrum antagonist of glutamine is more effective in inducing tumor regression than selective inhibition of a single glutamine-utilizing enzyme (36), DON is being re-evaluated as a potential therapy in a number of cancers specifically in combination with other chemotherapeutic agents (37, 38). Therapeutic strategies using DON against glutamine dependent tumors have also been proposed (38).

Owing to the central role of hexosamine biosynthesis pathway in pancreatic cancer (that is known to be heavily dependent on glutamine metabolism (39)), we evaluated DON in pancreatic cancer as both an anti-tumor and anti-stromal agent in the current study. Our results showed that DON acted as a potent anti-tumor agent and inhibited self-renewal and clonogenicity in pancreatic cancer cells. It also significantly decreased metastatic potential of pancreatic cancer cells. Further, when co-implanted with pancreatic cancer-associated fibroblasts in the pancreas of C57BL/6 mice, DON had a profound effect on the ECM and promoted infiltration of CD8+ cytotoxic T-cells. Further, infiltration of cytotoxic T-cells in pancreatic tumors following DON administration also sensitized them to anti-PD1 therapy. Since pancreatic tumors are notoriously immune evasive, this observation is extremely promising as it indicates that metabolic inhibitors like DON can be developed to overcome immune resistance and improve survival rates in this disease

## Results

### Hexosamine Biosynthesis Pathway is overactivated in pancreatic cancer

Since chronic pancreatitis is a well-known risk factor for pancreatic ductal adenocarcinoma(47), we studied the expression of GFAT1/2 (alias GFPT1/2) as well as the other enzymes in this pathway in caerulein-induced chronic pancreatitis as well as in KPC pancreatic cancer mouse model during tumor progression. Our results showed that these enzymes were overexpressed upon induction of pancreatitis (Figure 1A) as well as during pancreatic tumor progression (Figure 1B). In addition, expression of GFAT1 was also increased in the ductal cells of the pancreatic adenocarcinoma when observed in a tumor tissue microarray (Figure 1C). Further, an analysis of The Cancer Genome Atlas (TCGA) database showed that this pathway was overexpressed in 35.7% of the 176 pancreatic cancer patients in the database at both the RNA and protein level (Figure 1D). To study if GFAT1 was expressed both in the tumor and stroma, we performed immunohistochemistry with anti-alpha SMA and anti-GFAT1 antibody. Our results showed that GFAT1 is predominantly expressed in the tumor cells. As seen in the figure (Figure 1E, F), GFAT1 did not co-stain with alpha SMA in the mouse KPC tumors (Figure 1E) or in the human tumors (Figure 1F). Since GFAT1 is the rate-limiting step of this pathway, we focused our study on this particular enzyme.

**Figure 1.**
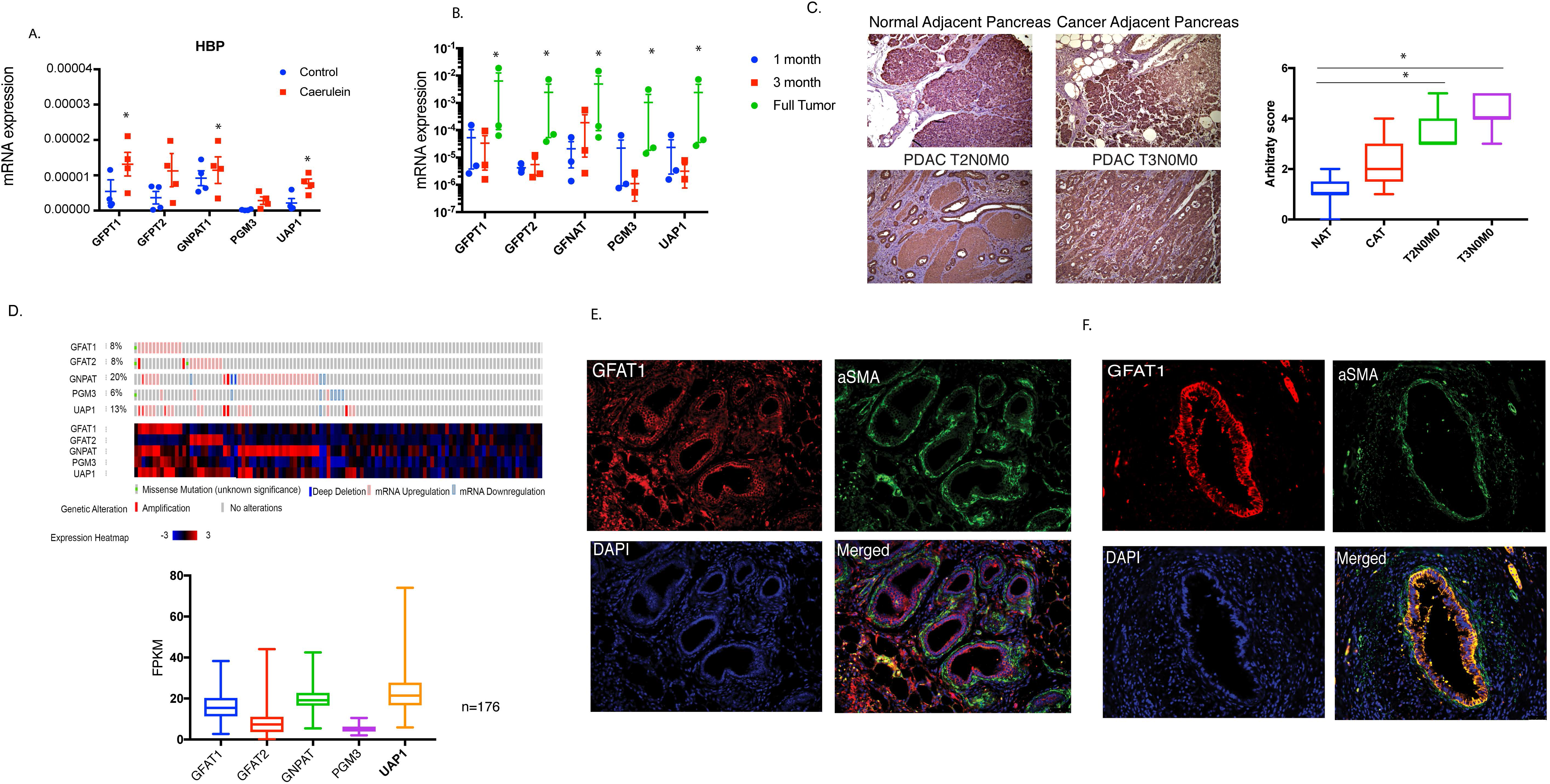
Hexoamine Biosynthesis Pathway is activated in PDAC and chronic pancreatitis. Enzymes in the hexosamine biosynthesis pathway are overexpressed in pancreatitis (A) as well as in pancreatic cancer mouse model KRas^G12D^TP53^R172H^-Pdx^Cre^,or KPC.The expression of enzymes increased as the tumor progressed (B). In tumor tissue microarray of PDAC patients, GFAT1 (magnification 20X), the rate limiting step of hexosamine biosynthesis pathway was overexpressed. FPKM refers to Fragments Per Kilobase of transcript per Million mapped reads that correlates with relative expression of a transcript proportional to the number of cDNA fragments that originate from it. The increased expression correlated with advanced grade of the tumor (C).The microarray contained 2-3 samples of each disease stage. According to http://www.cbioportal.org, a large number of patient cohorts in TCGA showed alterations in the genes of hexosamine biosynthesis pathway (n=176) (D). GFAT1 expression (magnification 20X) was predominantly in the ductal cells as seen in tumors from KPC mice (E) or patient tumor tissue (F).The picture is representative of 3 patient samples and 10 fields per sample. All gene expression studies with qPCR were done 3 independent biological replicates. The error bars represent Mean +/-SEM. * represents p-value <0.05.

### GFAT1 contributed to aggressive biology of pancreatic cancer by regulating self-renewal and metastasis

A mark of an aggressive tumor is its ability to metastasize and its potential to relapse after treatment. These are dependent on the genes that regulate self-renewal. Our previous results (32) show that OGT, an enzyme dependent on UDP-GlcNAc and thus HBP, was instrumental in regulating self-renewal in pancreatic cancer via its effect on Sox2. Our results showed that inhibition of GFAT1, the rate-limiting enzyme of HBP, using siRNA resulted in inhibition of a number of self-renewal genes like Sox2, Oct4 and KLF4 in the pancreatic cancer cell lines MIA-PaCa2 and S2VP10 (Figure 2A). Since GFAT1 activity is dependent on availability of glutamine, we next blocked glutamine utilization with DON. Our studies showed that treatment of pancreatic cancer cell lines MIA-PaCa2 and S2VP10 with DON resulted in decreased expression of self-renewal genes as seen with GFAT siRNA (Supplementary Figure 1A). To study if the inhibition of HBP by blocking glutamine utilization with DON resulted in decreased clonogenicity (a surrogate assay for self-renewal), we performed a colony forming assay on the pancreatic cancer cell line S2VP10, which is aggressive and has high self-renewal capability. Our results showed that treatment with DON resulted in decreased colony formation showing that glutamine utilization by HBP was instrumental in decreasing self-renewal in pancreatic cancer cells (Figure 2B). This observation was further validated in the pancreatic cancer cell line L3.6PL (Supplementary Figure 1B). These observations indicated that DON suppressed self-renewal ability of pancreatic cancer cells. Our previously published data show that DON affected tumor cell proliferation (33). Our current study showed that treatment with DON decreased viability of primary KPC cells while it did not have any effect on the viability of primary cancer-associated fibroblasts in vitro (Figure 2C), indicating that within a tumor DON had differential effect on the cellular components.

**Figure 2.**
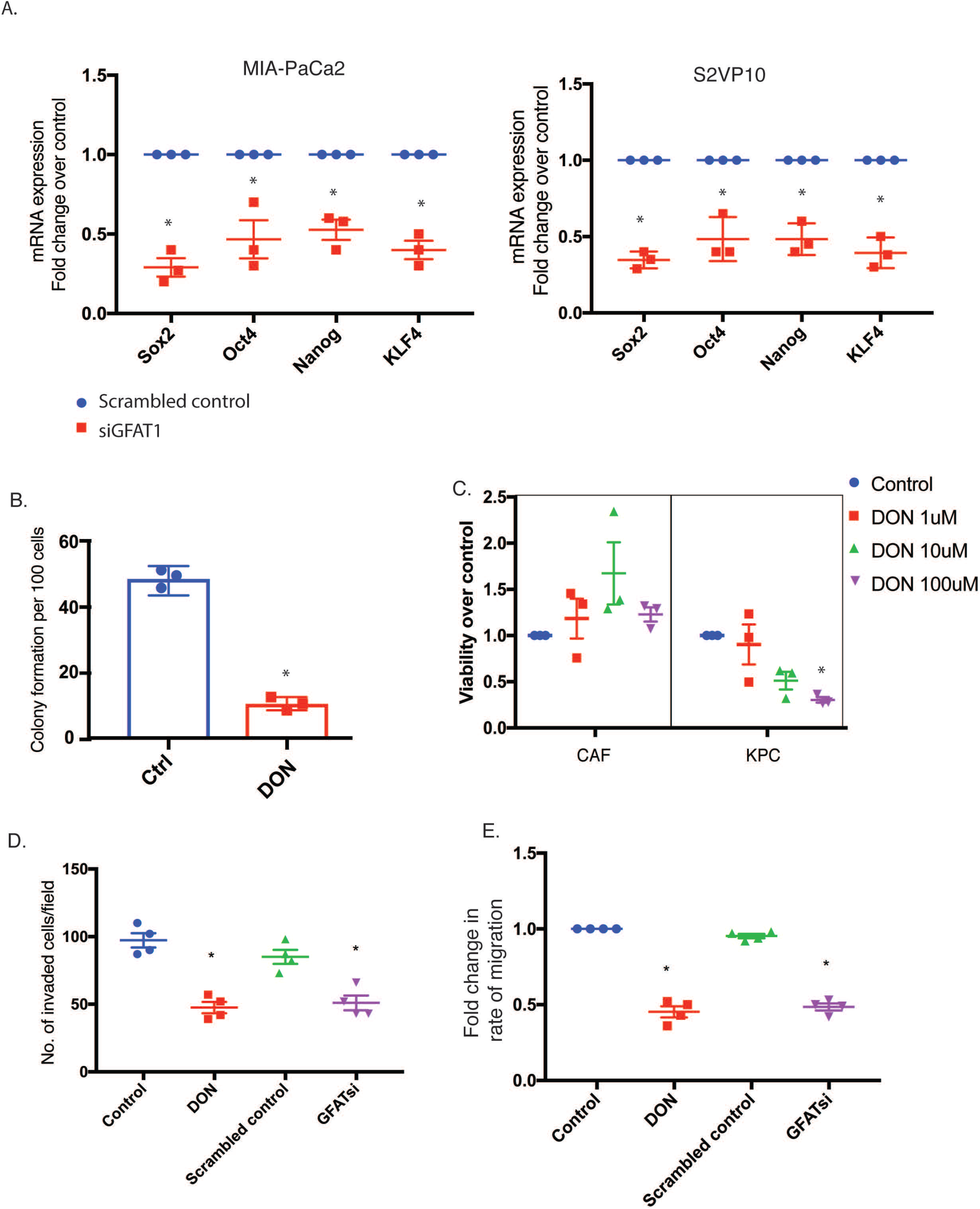
GFAT1 regulates self-renewal and invasion in PDAC. GFAT1 inhibition by siRNA decreased expression of self-renewal genes in pancreatic cancer cell lines MIA-PACA2 and S2VP10 (A). Treatment with glutamine analog DON at 50uM decreased colony formation in S2VP10 cells (B), indicating a loss of clonogenicity. Tumor epithelial cells (KPC) were more vulnerable to treatment with DON compared CAF cells (C). LC50 of DON for KPC cells was calculated to be 72μM. At non-lethal dose of 50µM, treatment with DON decreased invasion (D) as evaluated by Boyden Chamber assay, as well as migration of pancreatic cancer cells S2VP10 as tested by Electric Cell-Surface Impedance Sensing or ECIS (E). All gene expression studies with qPCR were done 3 independent biological replicates.Students t test (parametric) was used for statistical analysis and the data represents mean <u>+/− SEM.</u>

Since HBP affects the activity of OGT, which in turn is instrumental in regulating the metastatic property of cancer cells (48–50), we next evaluated the effect of blocking HBP on invasion and migration. Treatment with DON as well as GFAT siRNA decreased the invasiveness of S2VP10 cells when evaluated in a Boyden chamber assay (Figure 2D). Treatment with DON further decreased migration of pancreatic cancer cell lines when evaluated real-time via Electric Cell Surface Impedance Sensing or ECIS (Figure 2E), further showing that DON suppressed invasiveness and metastatic potential of aggressive pancreatic cancer cells.

### Inhibition of GFAT1 or glutamine utilization by DON resulted in regressed tumors and decreased metastasis in animals

To study if DON was efficacious in vivo, we implanted metastatic pancreatic cell line S2VP10 subcutaneously in the flanks of athymic nude mice. Treatment with DON (1mg/kg) decreased tumor progression (Figure 3A) as well as end-of-study tumor weight and volume (Figure 3 B, C). In addition, treatment with DON decreased Ki-67+ cells in the tumor indicating a loss in proliferative pancreatic cancer cells (Figure 3D, Supplementary Figure 2A).

**Figure 3.**
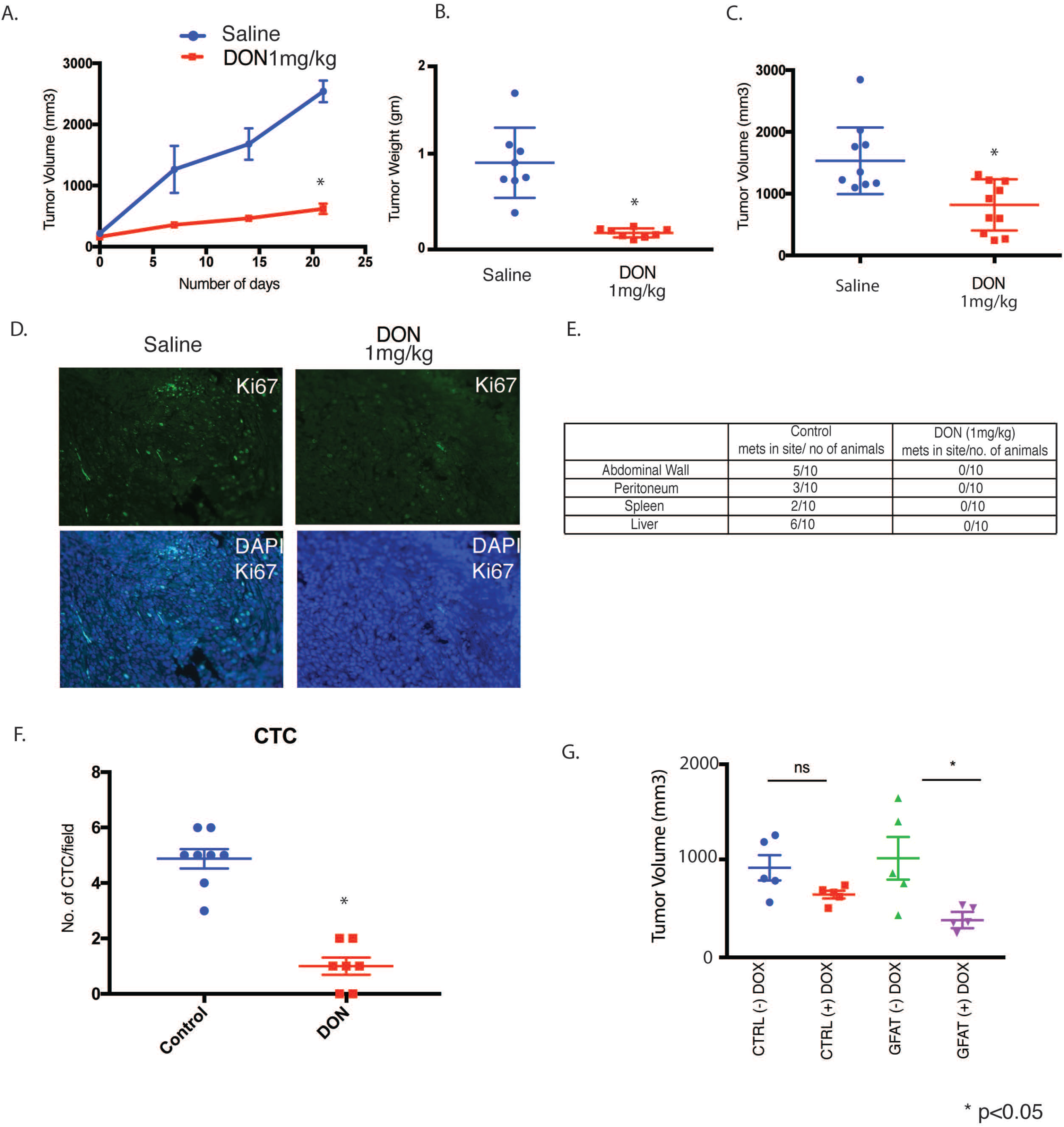
Inhibition of GFAT1 has anti-tumor effect in vivo. Treatment with 1mg/kg/5 days a week DON decreased tumor progression in subcutaneous implantation of pancreatic cancer cells S2VP10 in athymic nude mice (n=10) (A). Endpoint tumor weight (B) and volume (C) were significantly decreased following treatment. Ki67 staining showed decrease in the number of proliferating cells (D). Orthotopically implanted KPC:CAF in the pancreas of C57BL6 were treated with DON (1mg/kg/3 days a week) for 4 weeks. Treatment group showed decreased metastatic spread (E). Consistent with this, DON treated animals had less circulating tumor cells in their bloodstream (F). To confirm that anti-tumor effects were due to GFAT1/2 inhibition by DON, tet-inducible shGFAT1 cell-line was developed and implanted orthotopically in athymic nude mice. As observed with DON, upon induction of shGFAT1 with doxycycline, the tumor volumes of the animals were significantly smaller compared to the no doxycycline tumors (G).Students t-test, non-parametric (Mann Whitney test) was used to determine statistical significance and data represents mean <u>+/− SEM. N= 10 mice was used for all groups.</u>

To test the efficacy of inhibition of glutamine utilization on metastasis, we implanted KPC and CAF cells orthotopically in the pancreas of C57BL/6 mice at a ratio of 1:9. DON (1mg/kg/3 days a week) was administered for 30 days. Metastatic spread to local and distant tissues was documented following necropsy. DON significantly decreased metastatic spread and increased necrosis (Figure 3E, Supplementary Figure 2B). In addition, DON significantly decreased the number of circulating tumor cells in this model (Figure 3F, supplementary Figure 2C).

To rule out off-target effects of small molecule DON, we next constructed a tet-driven shGFAT1 and transfected metastatic pancreatic cancer cell line S2VP10 to generate a tet-shGFAT1 cell line. These cells were implanted orthotopically in athymic nude mice. Ten days after implantation, GFAT1 expression was turned off by adding tetracycline to the chow. Animals were followed for an additional 30 days. As seen with DON, there was a significant reduction in end-of-study tumor volume (Figure 3G and of Collagen and HABP staining (Supplementary figure 2D, E)

### Treatment with DON modulates ECM in PDAC

Since HBP is responsible for synthesis of UDP-GlcNAc, which is also the substrate for hyaluronan (HA), a major extracellular matrix component in pancreatic tumors, we next evaluated the effect of DON on ECM components of pancreatic tumors. To study this, KPC001 and CAF cells (1:9 ratio) were implanted in the pancreas of C57BL/6 mice and treated with DON(1mg/kg/3 days a week). Our studies showed that animals treated with DON had lower HA (Figure 4 A Supplementary Figure 3A) as well as lower collagen (Figure 4B, Supplementary Figure 3B). To study if this decrease in HA and collagen was due to decreased gene transcription, we next studied the expression of genes involved in their synthesis. Our study showed that expression of HA synthase 1 (HAS1) by the KPC tumor cells was significantly downregulated in the DON-treated group (Figure 4C). Since other ECM components are equally synthesized by tumor epithelial cells and stromal fibroblasts, we next co-cultured KPC001 (murine primary tumor cells) and CAF cells and treated them as indicated in Supplementary Figure 3C. Our study showed that while there was a significant decrease in a number of collagen synthesis genes along with expression of genes involved in other structural components of ECM (Figure 4D), there was a more profound effect on the ECM remodeling proteases (Figure 4E). This indicated that there was an extensive ECM remodeling in the pancreatic tumor microenvironment. In addition, there was also a significant alteration of expression of a number of cell-cell and cell-ECM adhesion molecules (Supplementary Figure 3D).

**Figure 4.**
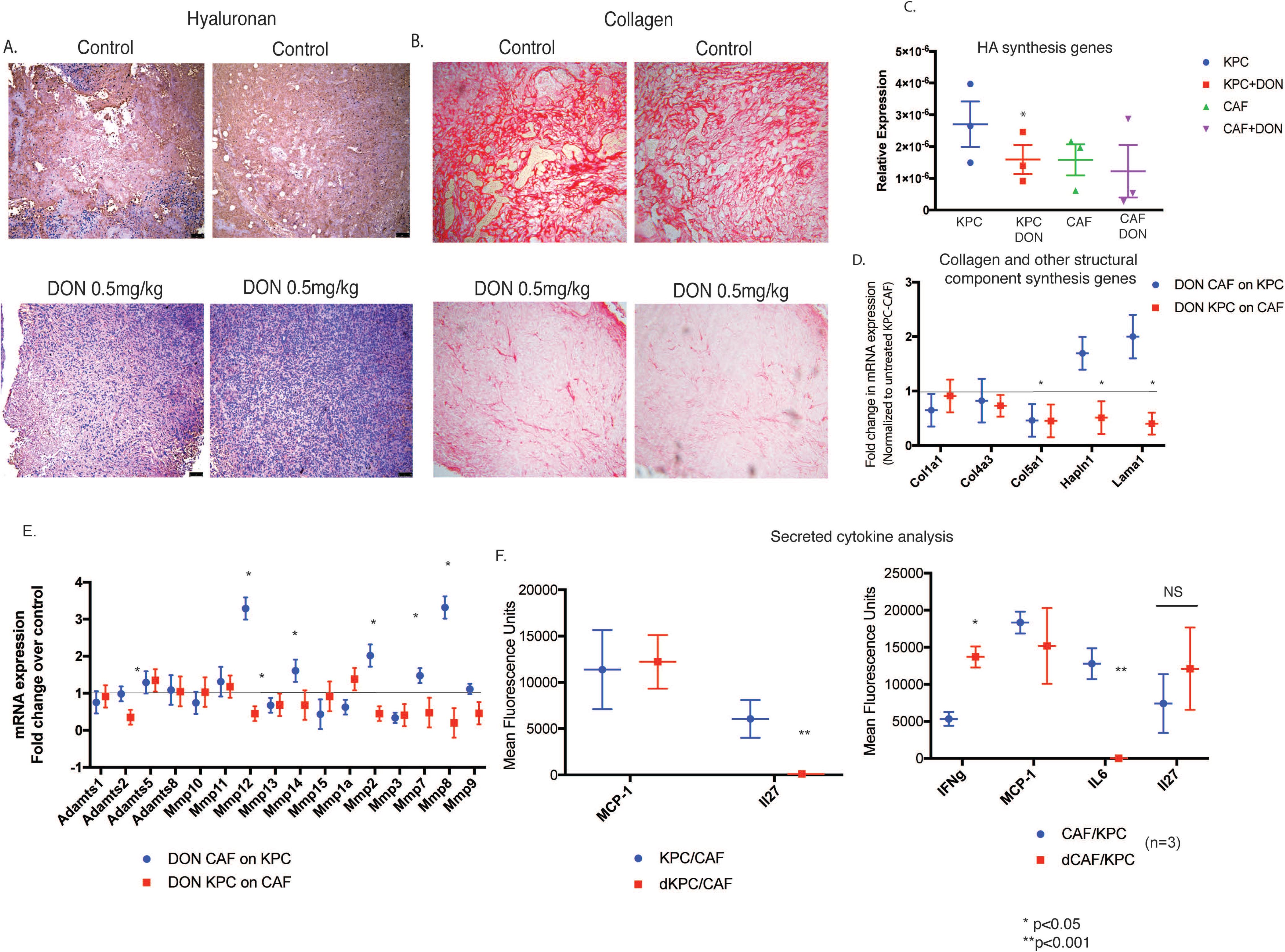
Treatment with DON remodeled ECM in orthotopic syngeneic mouse model. KPC and CAF cells were implanted orthotopically in the pancreas of C57BL6 mice and treated with 0.5mg/kg DON/ 3 days a week for 30 days. Treatment with DON significantly decreased hyaluronan (A) and collagen (B) in the extracellular matrix of the tumor. Genes responsible for hyaluronan synthesis, such as (HAS1), were decreased significantly in the KPC cells but did not change in CAF cells (C). In a co-culture experiment, treatment with DON (50µM) significantly decreased collagen synthesis genes in both KPC and CAFs, while other structural component genes like Hapln1 and Lama1 were only decreased when DON treated KPCs were co-cultured with CAFs (D). Treatment with DON also changes expression of a number of ECM proteases significantly (E). Further, the secreted IL27 was significantly decreased in DON treated KPC cells, while IL6 was significantly decreased in DON treated CAF cells. Secretion of IFNγ was increased in CAFs upon treatment with DON (F).Each experiment was repeated 3 times and the result is represented as mean <u>+/− SEM. Students t test was used for determining statistical significance.</u>

It is well-known that the tumor and stromal cells remodel the ECM not only via synthesis of basement membrane constituents, but also through cytokine secretion. To evaluate this, we next set up the co-culture as described in Supplementary Figure 3C, and estimated the secreted cytokine profile using a cytometric bead array. Upon co-culture in a transwell, in which CAFs were treated with DON, IFNγ and IL6 were observed to be significantly altered by DON treatment, while only IL27 showed a significant decrease when only KPC cells were treated with DON in the co-culture (Figure 4F). Treatment of tumor cells (KPC) with DON completely abolished the secretion of IL27 in the co-culture, while treatment of CAF cells with DON did not significantly change the secretion of this cytokine. In CAFs, IL6 secretion was significantly inhibited upon treatment while changes in MCP1 and IL27 were not significant. Since IL6 and IL27 are both pro-tumor cytokines, its downregulation by DON indicated a profound anti-tumor activity of this compound. Similarly, IFNγ plays a role in activation of M1 macrophages as well as infiltration of T-cells in the tumor, eliciting a tumor tissue disruptive effect (51). These changes following treatment with DON indicated that DON played an anti-tumor role by affecting the tumor microenvironment of the pancreatic tumor (52).

### Inhibition of GFAT1/HBP affects immune landscape in PDAC

It is well known that remodeling the ECM in a tumor affects its immune landscape (22). Based on the change in the secretion profile of cytokines from the tumor cells and stromal cells upon treatment with DON, it seemed likely that this would significantly affect the infiltration and function of immune cells in the pancreatic tumor. Macrophage density has been correlated with overall survival in pancreatic cancer patients (53). Our results showed that treatment with DON resulted in an increase in the activated macrophage population as seen by CD68 staining (Figure 5A, B). Since our previous results showed that treatment of pancreatic tumor cells with DON (KPCs or CAFs) increased IFNγ (Figure 4F), it is possible that the increased CD68+ macrophage population within DON-treated tumors is a direct consequence of that event.

**Figure 5.**
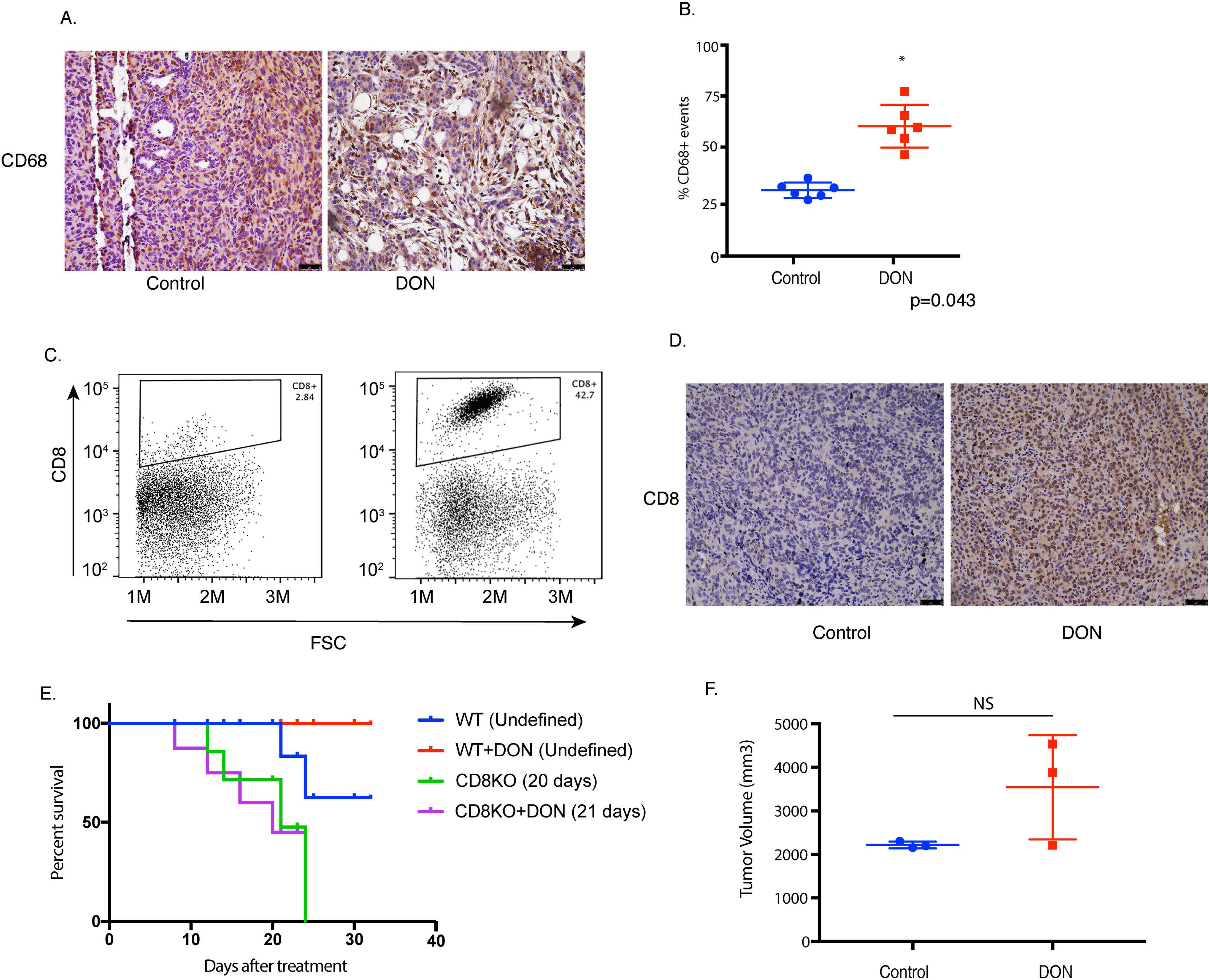
Treatment with DON altered immune profile in orthotopic syngeneic mouse model: In the orthotopic implantation of KPC and CAF in the pancreas of C57BL6 mice, DON increased CD68+ macrophage population (A, B). Treatment with DON also increased intra-tumoral CD8+ infiltration as seen by flow cytometry (C) and immunohistochemistry (D). Effect of DON on tumor and stroma in syngeneic model was CD8 mediated as pancreatic tumors implanted in CD8KO mice did not show improved survival (E) or decrease in tumor volume (n=3) (F) with DON.The image for CD8 was acquired at 10X magnification.

Our analysis of the tumor tissue further showed that there was a significant increase in the intra-tumoral CD8+ T-cells (Figure 5C, D). Increased infiltration of T-cells is associated with better prognosis in pancreatic cancer. This indicated that DON was modulating the immune milieu of the pancreatic tumors by increasing the CD68+ macrophages and promoting increased infiltration of CD8+ T-cells, thereby reversing the immune suppressive microenvironment.

To study if DON was still an effective anti-tumor compound in the absence of CD8+ T-cells, we next implanted KPC001 and CAF cells orthotopically in the pancreas of CD8KO mice in a ratio of 1:9. The tumors were allowed to grow for two weeks before they were randomized to DON (0.5mg/kg/3 days a week) treatment or control groups. Treatment was continued for 30 days and survival analysis was performed. Our results showed that survival of DON-treated animals in the CD8KO mice was not different from the untreated animals (Figure 5E). Further, necropsy of animals across both groups showed no difference in tumor volume (Figure 5F). These results suggested that efficacy of DON in an immune competent syngeneic pancreatic cancer model was largely dependent on CD8+ T-cells.

### Inhibition of GFAT1/Glutamine utilization sensitizes to anti-PD1

Increased CD8+ T-cells within a tumor has been associated with increased sensitivity to immune therapy in a number of cancers (54). Since treatment with DON alone resulted in modulating the ECM, altered cytokine secretion, increased activated CD68+ macrophages as well as cytotoxic CD8+ T-cell infiltration in the pancreatic tumor, we next evaluated whether treatment with DON also made pancreatic cancer susceptible and sensitive to immune therapy. To study this, KPC001 and CAF cells were implanted orthotopically in the pancreas of C57BL6 mice in a ratio if 1:9. The tumors were allowed to grow for 14 days following which animals were randomized into 4 groups: Control/Isotype Ab, DON (0.5mg/kg), anti-PD1 Ab (100ug/3 injections), anti-PD1 + DON (0.5mg/kg). Treatment was continued for 1 month after which the animals were sacrificed. Our results showed a profound effect on the tumor weight (Figure 6A) and tumor volume (Figure 6B) with the combination of DON and anti-PD1. Additionally, the combination of DON and anti-PD1 resulted in better survival advantage compared to either group alone (Figure 6C). Further, the combination resulted in a decrease of PDL1 expression in the tumor, indicating that this immune evasive property of the tumor was being overcome by DON (Figure 6D). Assessment of the ECM components following treatment with DON and anti-PD1 antibody showed that DON alone as well as in combination with anti-PD1 decreased HA and collagen (Figure 6E, Supplementary Figure 4 C,D,E). Additionally, we also analyzed the expression of other checkpoint inhibitors in the tumor. Our results showed that while expression of B7-H3 did not change with DON, expression of TIM3 and CTLA-4 were down regulated following treatment with DON (Supplementary Figure 5A-C). We also analyzed the Immune cells from the spleen of tumor bearing mice. As seen from the tumor, the CD8+ cells were higher in DON treated mice. Further characterization showed these CD8+ cells to have low PD1 expression, showing that they were not exhausted (Supplementary Figure 5D-J).

**Figure 6.**
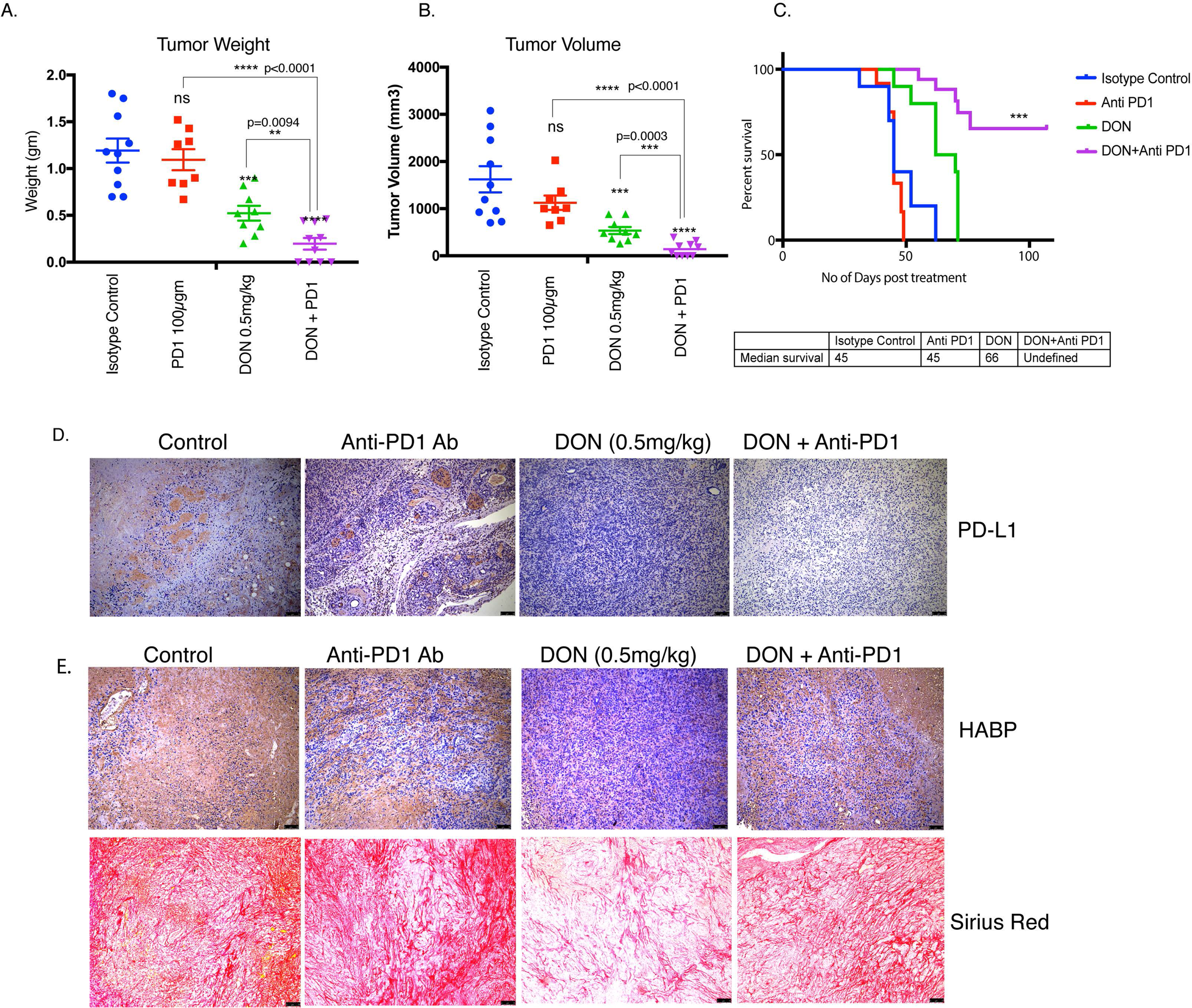
Treatment with DON sensitized PDAC to anti-PD1 antibody. Pancreatic tumor bearing mice were treated with Isotype Ab, DON (0.5mg/kg/ 3 days a week). Anti PD1 Ab(100μgm/3 times) and DON(0.5mg/kg/ 3 days a week) + Anti-PD1 Ab (100μgm/3 times). Treatment with DON as well as with DON+ anti-PD1 Ab showed significant decrease in tumor volume (A) as well as in tumor weight (B). Combination treatment also increased survival of tumor bearing animals (C) DON decreased PDL1 expression as seen by immunohistochemistry image (D). Treatment with DON decreased HA and collagen in the syngeneic tumors, however the decrease of these ECM components in the combination group was marginal (E).The histological images for PDL1,HA and Collagen were acquired at 10X magnification. 10 mice were used in each group and the result is represented as mean <u>+/− SEM. ANOVA was used for statistical analysis.</u>

## Discussion

Glutamine is one of the metabolites that is essential for growth and proliferation of cancer cells. It fuels a number of biosynthetic reactions in a cancer cell (55, 56). Consistent with this, glutamine-utilizing enzymes like GFAT1 are overexpressed in most cancers, including pancreatic cancer as reported by our lab (Figure 1) and others (57, 58). Pancreatic cancer cells have been shown to utilize glutamine in a non-canonical fashion to support proliferation via redox homeostasis (39, 59–62). As glutamine metabolism is dispensable for non-malignant cells, yet has a critical role in PDAC survival, it is an ideal candidate for therapeutic targeting.

Among glutamine-utilizing enzymes, glutamine-fructose amidotransferase 1 (GFAT1) is the rate-limiting enzyme for the hexosamine biosynthesis pathway (HBP). This pathway is a shunt pathway of glycolysis that utilizes fructose-6 phosphate from glycolysis and glutamine to synthesize UDP-GlcNAc, a nucleotide sugar that is essential for glycosylation reactions. Apart from N-glycosylation and O-GlcNacylation, UDP-GlcNAc is also a substrate for hyaluronan synthesis. Hyaluronan is a key component of the extracellular matrix and therefore, inhibition of HBP is likely to affect the extracellular matrix (ECM) in a tumor (63–65) (15). Thus, inhibition of GFAT1 effectively inhibits metabolic flux through this pathway and is likely to affect the glycosylation profile of the tumor cells as well as the components of the ECM. The ECM in a tumor actively drives tumor progression by providing a dynamic niche that regulates both mechanical as well as signaling abilities of a tumor cell (22). The ECM further plays a role in efficient infiltration of T-cells in the pancreatic tumor. Thus, targeting glutamine utilization is likely to remodel the pancreatic cancer microenvironment and make tumors amenable to immune therapy. This makes DON, a glutamine analog, very well suited for evaluation against PDAC as it is likely to have anti-tumor effect (by directly targeting signaling pathways through its effect on glycosylation reactions) as well as an anti-stromal effect (by interfering with HA synthesis in the ECM). Targeting just the microenvironment in pancreatic tumor, whether just cellular components (such as fibroblasts) or acellular components (like ECM), has been controversial (11, 66, 67). In fact, the stroma has been proposed to be a “restraining” mechanism of the host cells (13). However, preclinical studies from our lab as well as others have shown that the tumor microenvironment can be targeted only when the targeting agent can affect both tumor epithelial cells as well as stromal cells (10, 68, 69). Thus, treatment with DON, which increases anti-tumor activity and remodels the ECM to promote infiltration of cytotoxic T-cells in combination of anti-PD1 therapy, is a promising strategy to combat pancreatic cancer.

Treatment with DON primarily affects the ECM components of the tumor and as expected decreases the non-cellular components of the tumor. Our study showed that preventing utilization of glutamine (with DON treatment), and therefore inhibiting HBP, has a profound effect on the ECM composition of pancreatic tumors (Figure 4). Both collagen as well as hyaluronan (HA) in the ECM showed a decrease in the treated samples. Similarly, inhibition of GFAT in the tumors by tet-inducible shGFAT1, shows similar effects on collagen and HA content within the tumor (Supplementary Figure 2D). In addition, treatment of a KPC/CAF co-culture with DON *in vitro* showed that there was extensive downregulation in the expression of ECM proteases such as MMP12, MMP7 and MMP8 (Figure 4). These proteases are involved in dynamic remodelling of the ECM during tumor progression, specifically regulating metastasis (70). Recent studies have shown that amino acid mediated metabolic crosstalk between cancer associated fibroblasts and tumor epithelial cells affect the biophysical as well as biological properties of the tumor (71). Inhibition of HBP with DON (or by shGFAT1) resuls in altered expression of the ECM proteases along with the synthesis of genes involved in ECM-cell adhesion, indicating an extensive remodeling of the tumor microenvironment.

Treatment with DON also increases the infiltration of CD68+ macrophages (Figure 5A, B). This could be a consequence of increased IFNγ (Figure 4F) in the tumor cells. CD68+ macrophages are considered to be “anti-tumor” as they secrete tumoricidal cytokines like TNFα, IL12, reactive nitrogen species and oxygen intermediates. Additionally, CD68+ macrophages also promote infiltration of cytotoxic CD8+ T-cells (51). Our results show that DON indeed promotes infiltration of CD8+ T-cells in addition to an increasing CD68+ macrophages.

Apart from promoting tumor progression and metastasis, the ECM plays an active role in maintaining the immune landscape within a tumor. ECM regulates the migration of T-cells toward the tumor cells by providing a three-dimensional matrix as well as a chemokine gradient (22). Lack of infiltrating T-cells is considered to be one of prime reasons for the immune evasive phenotype of pancreatic cancer. Among the small population that does infiltrate, the robust ECM prevents it from migrating to the tumor cells in order to execute their cytotoxic activity. Our studies show that DON profoundly increases infiltration of CD8+ cytotoxic T-cells (Figure 5). *In vitro*, treatment with DON did not seem to affect T-cell viability, but significantly increases T-cell migration (Supplementary Figure 4B). Thus, remodeling of the ECM by DON actually promotes macrophage activation along with T-cell infiltration and migration into the pancreatic microenvironment. Our experiments further show that the effect of DON in the immune competent syngeneic mice was being mediated via increased infiltration of CD8+ T-cells as this was lost in tumors implanted in CD8 knock-out animals (Figure 5 E,F). In these animals, the tumor burden did not decrease following DON treatment. Our studies further show that DON sensitizes tumors to anti-PD1 therapy (Figure 6 A, B). The presence of CD8+ T-cells within the tumor improves response to anti-PD1 therapy (72). Thus, increased CD8+ T-cells as a result of treatment with DON sensitizes the pancreatic tumors to anti-PD1 therapy.

DON affects the tumor epithelial cells along with the ECM. Treatment with DON in a subcutaneous model of pancreatic cancer decreases tumor progression and also decreases tumor weight and volume (Figure 3). In an orthotopic model of pancreatic cancer, DON decreases metastatic spread of the tumor as well (Figure 3). This anti-metastatic property of DON is of significance, since DON also decreases the ECM production in pancreatic tumors. As the remodeled and reduced ECM would have normally promoted tumors to metastasize to distant organs, the anti-tumor activity of DON prevented that. However, even though tumor cells responded to DON treatment, the stromal fibroblasts (or CAFs) were resistant to DON treatment, as seen by the viability assay done in vitro (Figure 2C). This was an interesting observation since athymic nude mice are immunocompromised, and if DON was facilitating its effect solely by altering the immune landscape, we would not have seen such an astounding effect on tumor regression in these mice. However, DON also has a profound effect on cancer cells as it prevents glutamine utilization (since it is a glutamine analog). It is possible that in our *in vivo* experiment with the nude mice, where the tumors lack ECM and immune cells, the effects observed are purely due to the anti-tumor effect of DON. In the immune competent mice, where the tumors were co-implanted with CAFs that secrete a robust ECM, the ECM remodeling effects of DON promoted cytotoxic T-cell infiltration and sensitized to the anti-PD1 therapy. Taken together, this indicates that the combination affect of DON on inhibition of the cancer cells to metastasize and the ECM remodeling to allow immune cell infiltration is responsible for the profound anti-tumoral affect of DON treatment.

Converting “cold” pancreatic tumors that are unresponsive to immune therapy to a “hot” and responsive tumors is a focus of pancreatic cancer research. Our studies show that DON, a glutamine analog, can be used for this. The anti-tumor effect of DON was observed earlier during Phase 1 and 2 clinical trials in a number of cancers including lung and colon cancer. Data from multiple trials demonstrated that DON was safe and could potentially be used as a single agent. These studies were conducted in the 1980’s and in 53% of patients there were reports of a stable disease in colorectal cancer. Clinical trials with DON never reached Phase III trials. The dose for DON used in these studies ranged from 50mg/m^2^-480mg/m^2^. The lowest dose used (50mg/m^2^) translates to roughly 6mg/kg in mice (35, 38, 73). However, DON was never evaluated as a sensitizing agent for immune therapy in cancer. The potent effect of DON on ECM appeared to promote an increase in tumoricidal macrophages and an infiltration of cytotoxic T-cells in our study. This implied that targeting glutamine utilization with DON in pancreatic cancer can be developed as a potential therapeutic option to sensitize the normally immune resistant pancreatic cancer to anti-PD1 therapy.

## Conclusion

Pancreatic ductal adenocarcinoma (PDAC) is considered to be a highly immunosuppressive and heterogeneous neoplasm (6, 74). Despite improved knowledge regarding the genetic background of the tumor and better understanding of the tumor microenvironment, immune checkpoint inhibitor therapies (using CTLA-4, PD-1, PD-L1), that have shown effect in other solid tumors, have not been very successful in PDAC. Thus, novel therapeutic strategies that can make PDAC more amenable to immune therapy are urgently needed. In this context, our study is extremely timely as it reveals how specific metabolic pathways can be targeted in cancer cells to overcome immune resistance and sensitize the tumors to anti-PD1 like therapy options that have worked extremely well in other cancers.

## Materials and Methods

### Cell lines, treatments and reagents

SU.86.86(ATCC® CRL-1837) and MIAPaCa-2(ATCC® CRM-CRL-1420) were purchased from ATCC and were cultured according to the recommended conditions. Human Pancreatic Stellate Cells (3830) were purchased from ScienCell. Primary KPC cell line was isolated from a tumor of a 5-6 month old genetically engineered mouse model of KRAS^G12D^P53^R172H^Pdx-1-Cre (KPC) mice. The cells were isolated according to the protocol from our previous study (40) Cancer-associated fibroblasts (CAFs) were isolated from KPC mice according to the protocol described by Sharon et al (41). The purity of the fibroblasts was evaluated by flow cytometry using fibroblast surface protein (FSP) antibody and CK19 antibody. FSP+ and CK19-population was used for subsequent experiments. All the established cell lines were used from passages 5-20. S2-VP10 cells were a generous gift from Dr. Masato Yamamoto’s laboratory, University of Minnesota, Minneapolis, MN. Pancreatic Stellate Cells from mouse pancreas was isolated according to the protocol as described by Apte et al (42). SU.86.86 and S2-VP10 cells were grown in RPMI 1640 (Gibco) containing 10% FBS and 1% penicillin/streptomycin (Gibco). MIA PaCa-2, KPC and CAFs were grown in DMEM high glucose containing 10% FBS and 1% penicillin/streptomycin. All cell lines were routinely tested for mycoplasma and STR profiles (ATCC). 6-Diazo-5-oxo-L-norleucine, DON (D2141) was purchased from SIGMA ALDRICH and was used at a dose of 50μM for *in vitro* experiments.

### Colony forming assay

S2-VP10 cells were pretreated with DON for 24 hours. Cells were counted and plated at a density of 10K,1K and 100 cells per well in both treatment and control group respectively. Colonies were counted after 4 days of plating. The results represented are from 1000 cells per well but the result is similar across different dilutions.

### Immunohistochemistry

Tissues were deparaffanized by heating at 56°C overnight and then hydrated by treating with Xylene (15 mins, two times),100% Ethanol, 90% Ethanol, 70% Ethanol (2 times) 5 mins each. The slides were then steamed with a pH 6 reveal decloaker (Biocare Medical, Concord,CA) for antigen retrieval, blocked in Dako serum blocker (Agilent technology). Primary antibody was added overnight. Slides were washed 3X in PBS, secondary antibodies (Alexaflour) were diluted in SNIPER (BIOCARE Medical) and slides were stained for 30 mins at room temperature/ Slides were then washed again 3X in PBS and mounted using Prolong Gold anti-fade with DAPI (Molecular Probe) for immunofluorescence staining. For Immunohistochemistry, the cells were then treated with DAB (SK-4100) and hematoxylin (H-3404,Vector Labs, Burlingame CA. The stained slides were then dehydrated by dipping in 70% Ethanol (2 times), 90% Ethanol, 100% Ethanol and Xylene (15 mins, two times). Slides were then mounted with paramount and dried overnight and imaged by a bright field microscope. GFPT1 antibody (14132-1-AP) was purchased from protein tech and used at a dilution of 1:200. α-SMA (ab5694), CD4 (ab183685), CD8 (ab22378) and PDL1 (ab80276) was purchased from Abcam and were used at a dilution of 1:200,1:1000,1:200 and 1:50 respectively. For CD4 and CD8 antibody citrate buffer was used for antigen retrieval. Ki-67 SP6 antibody was purchased from Thermo Fisher Scientific and was used at a dilution of 1:200.

### Animal studies

All experiments were done according to the protocol approved by Institutional Animal Care and Use Committee (IACUC) at University of Miami. Female athymic nude (nu/nu) mice between the ages of 4-6 weeks were used for in vivo experiments. For subcutaneous experiments 500,000 S2VP10 cells were implanted in the right flank of athymic nude mice. Corning® Matrigel® Growth Factor Reduced (GFR) Basement Membrane Matrix and 1X PBS at a ratio of 1:1 were used as a suspension medium for the cells. Tumors were allowed to reach a size of 100mm3 and then the treatment was started. Mice were given DON at a dose of 1mg/kg/5days a week for 4 weeks. At the end of the 4^th^ week mice were sacrificed and the tumor weight and volume were noted. Tissues were flash frozen for further experiments.

Female C57/BL6 mice between the ages of 4-6 weeks were used for orthotopic implantation of KPC and CAF cells. 1000KPC and 9000 CAF’s implanted orthotopically in 100% Corning® Matrigel® Growth Factor Reduced (GFR) Basement Membrane Matrix. Tumors were allowed to grow for 2 weeks after which the mice were randomized and treatment was started. DON was given at a dose of 1mg/kg/3days a week i.p and 2 DG was given at a dose of 1000mg/kg/day i.p. for 4 weeks after which the mice were sacrificed. Orthotopic tumors were harvested, dimensions were noted and metastasis was evaluated. Blood for CTC isolation was collected via cardiac puncture and blood from 3 mice per group was pooled for CTC.

For evaluating the combinatorial effect of DON and PD1 on tumor growth female C57/BL6 mice between the ages of 4-6 weeks were used. 1000KPC and 9000 CAF’s implanted orthotopically in 100% Corning® Matrigel® Growth Factor Reduced (GFR) Basement Membrane Matrix. Tumors were allowed to grow for 2 weeks after which the mice were randomized and treatment was started. DON was given at a dose of 0.5mg/kg/3days a week and PD1 was given at a dose of 100μgm per mice at day 15,17 and 19 after starting DON treatment. DON was given for 4 weeks after which the mice were sacrificed and tumor was harvested and metastasis was evaluated.

CD-8 knockout mice (B6.129S2-Cd8atm1Mak/J) were purchased from Jackson Laboratory. The mice are referred as CD8KO in this study.1000 KPC and 9000 CAF cells were implanted orthotopically in 100% Corning Matrigel Growth Factor Reduces Basement Membrane Matrix in female CD-8KO mice between 4-6 weeks of age. Tumors were allowed to form for 2 weeks after which the mice were randomized into control and treatment group. DON was started at week 3 post-surgery at a dose of 0.5mg/kg 3 times a week. Mice were monitored daily for morbidity or mortality and a survival curve was plotted.

### Isolation of Circulating Tumor Cells

Circulating tumor cells were isolated according to (43). Briefly, 0.5 ml of blood was collected from animals in the control and DON treated group in EDTA coated tubes. The blood was passed through the Circulogix FaCTChecker system and captured on the system’s filter cartridges mounted on slides. Captured cells were stained with fluorescent tagged antibodies against CD45 and CK19. The CD45-CK19 + fraction was quantitated from each group of animals.

### Multiplexed cytometric bead array

KPC and CAF cells were pre-treated with DON for 24 hours. 100,000 cells were plated in Corning® Transwell® polycarbonate membrane cell culture inserts in the following manner-Cells placed in the chamber have the letter “C” after them and the cells plated on the plate have the letter “P” after them. The treatment groups were KPC(C)+CAF(P), DON KPC(C)+CAF(P), DON CAF(C)+KPC(P) DON KPC (C)+DON CAF (P) and media only. Cells were allowed to be in coculture for 24 hours. Conditioned Media was collected post incubation. Mouse Inflammation Kit (BD 552364) was used and the experiment was performed according to the manufacturer’s instructions.

### Flow-cytometry for infiltrated immune cells

Tumor samples harvested from mice were placed in RPMI until they were ready to be processed. Tumors were minced into tiny pieces and then they were digested with Collagenase IV at 37 degree for 1-3 hours. After the tissue has been digested they were passed through a 40μm nylon filter. The tissue was then spun at 500g for 5 mins and was then resuspended in 1ml of flow buffer (0.5% Bovine Serum Albumin,2mM EDTA, 1% Penicillin-Streptomycin in 500mL PBS) The tubes were then spun again at 500g for 5 mins. 100μl of cytofix-cytoperm buffer was added to the pellet and the cells were allowed to fix for an hour at room temperature. After 1 hour, the cells were then washed with 1 ml of FACS buffer and then centrifuged at 500g for 5 mins. The supernatant was then discarded and the cells were stained with the surface and intercellular antibody for 40 mins in the dark. After 40 mins of staining, the cells were washed with 1ml of FACS buffer and then spun at 500g for 5 mins. Cells were then resuspended in 200μl of FACS buffer and were acquired. The antibodies used for T cell analysis were all from biolegend-CD3 PE Dazzle (100348), CD4 PE Cy7(10028), CD89AF-657), CD49b FITC (108906), CD25 AF-700 (102024),TCRγ/δ BV 510(118131), FoxP3 PE (126404),IL4 BV-711 (504133),IL-17 BV-421 (506926),TNF-a BV-650 (506333),IL10 APC/Cy7 (505036) and IFN-γ.

### Boydon Chamber Invasion Assay

24 well chamber inserts (Corning BioCoat) were hydrated in serum-free medium for 2-3 hours: the bottom of the chamber wells contrainted the attractant (10% FBS contraining DMEM) 50,000 cells were plated on the top of the insert. 24 hours later, cells that did not invade were scrubbed from the top chamber via a cotton swab and the invaded cells were fixed in methanol and stained with crystal violet. Invases cells were counted my microscopy.

### Migration Assay

The migration assay was conducted by Electric Cell-substrate Impedence Sensing (ECIS) (Applied Biophysics) and performed as described in Banerjee et al, 2014(44).

### Tetracycline inducible GFAT knockdown cell line

For the generation of tet inducible GFAT knockdown cell line. S2VP10 was selected as a parent cell line. shRNA oligoes for GFPT1 was purchased from idt and the cell lines were made according to the protocol as described by Wiederschain, D et al (45) and Wee et al(46)

### Statistics

All *in vitro* experiments were performed in 3 independent runs and the values were expressed as the mean ± SEM. Students t tests were used to determine the significance and P values of less than 0.05 were considered statistically significant.

### Study Approval

All animal studies were performed according to the protocol approved by the University of Miami Department of Veterinary Research (DVR) and Institutional Animal Care and Use Committee (IACUC). Deidentified Patient TMAs were either obtained from US Biomax Inc commercially.

## Author Contributions

Performed experiments, analyzed data, prepared manuscript: NSS,VKD,VTG,RH,BG,AF

Analyzed data, performed quantitation, revised manuscript: KK, BCD,VD

Conceptualized and supervised project, wrote and revised manuscript: SB

Provided resources, revised manuscript: VD, AS, SB

## Acknowledgements

The authors would like to acknowledge Oliver Umland at the Diabetes Research Intstitute, University of Miami for his support with flow cytometry.

