## Supplemental data for "Targeting tumor-intrinsic metabolic node sensitizes pancreatic cancer to anti-PD1 therapy"

### Supplementary Methods

#### T-cell migration assay

T-cells were freshly isolated from spleens of 6-8 week old C57/Bl6 mice using Mojosort T-cell isolation kit (Biolegend) according to manufacturer's protocol. Following isolation, they were activated by the Miltenyl T cell Activation/Expansion kit 9130-093-627) was used and the experiment was performed according to the manufacturer's instructions. The activated T cells were then plated in a 6-well plate and then treated with 50uM DON for 24 hours. The DON pre-treated T cells were then placed in a migration chamber. In the bottom chamber of a 6-well plate 100,000 KPC and CAF cells were placed at a ratio of 1:1 and the DON treated Activated T cells were placed on the top. The T cells were plated in DMEM containing 1%FBS and 1%Pen/Strep whereas the KPC/CAF cells in the bottom chamber were plated in DMEM containing 10% FBS and 1% Pen/Strep thereby creating a concentration gradient. The cells were allowed to migrate for 24 hours and then the chambers were fixed with Methanol and stained using 0.1% crystal violet.

#### Quantitation of images

All images were quantitated in a blinded manner by 3 independent research personnel in the laboratory. 3 slides from each group were imaged and at least 10 fields were captured from each slide. An arbitrary scoring unit was defined (0= no staining 4= maximum staining). The values obtained were averaged and represented in the manuscript.

Supplementary Figure Legends

Supplementary Figure 1. Treatment of pancreatic cancer cell lines MIA-PaCa2 and S2VP10 with DON resulted in decreased expression of self-renewal genes as seen with GFAT siRNA (A). DON decreased colony formation in pancreatic cancer cell line L3.6PL (B).

Supplementary Figure 2. Quantification of Ki67 stating in DON treated tumors (A). Quantification of necrosis (B), Image showing captured circulating tumor cells (C). Tet-inducible shGFAT1 showed a profound effect on HA and collagen content in the tumor (D,E).

Supplementary Figure 3. Quantitation of HA (A) and collagen (B) in the DON treated tumors. Schematic diagram showing set-up of co-culture experiments with KPC and CAF cells for ECM analysis and cytokine profile analysis (C). mRNA expression of ECM genes deregulated upon treatment with DON.

Supplementary Figure 4. Representative image showing migration of T-cells following DON treatment (A). DON treatment did not change the viability of T-cells, but it increased T-cell migration (B). Quantitation of PD1 staining in the tumors treated with DON, Anti PD1 antibody and combination of the DON and anti-PD1 (C). Quantitation of HA (D) and collagen (E) in the treated tumors.

Supplementary Figure 5. mRNA expression of DON treated tumors for checkpoint markers B7-H3 (A), TIM3 (B) and CTLA4 (C). Analysis of tumors showed that the infiltrated CD8 T-cells were mostly activated (D). DON did not change the proliferation (E) or PD1 expression (F) of these infiltrated T-cells. Upon analysis of the spleen, we observed an increased CD8+ T-cells in the spleen of pancreatic tumor bearing animals (G), that showed no change in proliferation (H). DON did decrease the PD1 expression in these cells, indicating that they were not exhausted (I). Most of the CD8+ cells in the spleen were naïve as seen by CD62L staining and not activated as seen by CD44 staining (J).

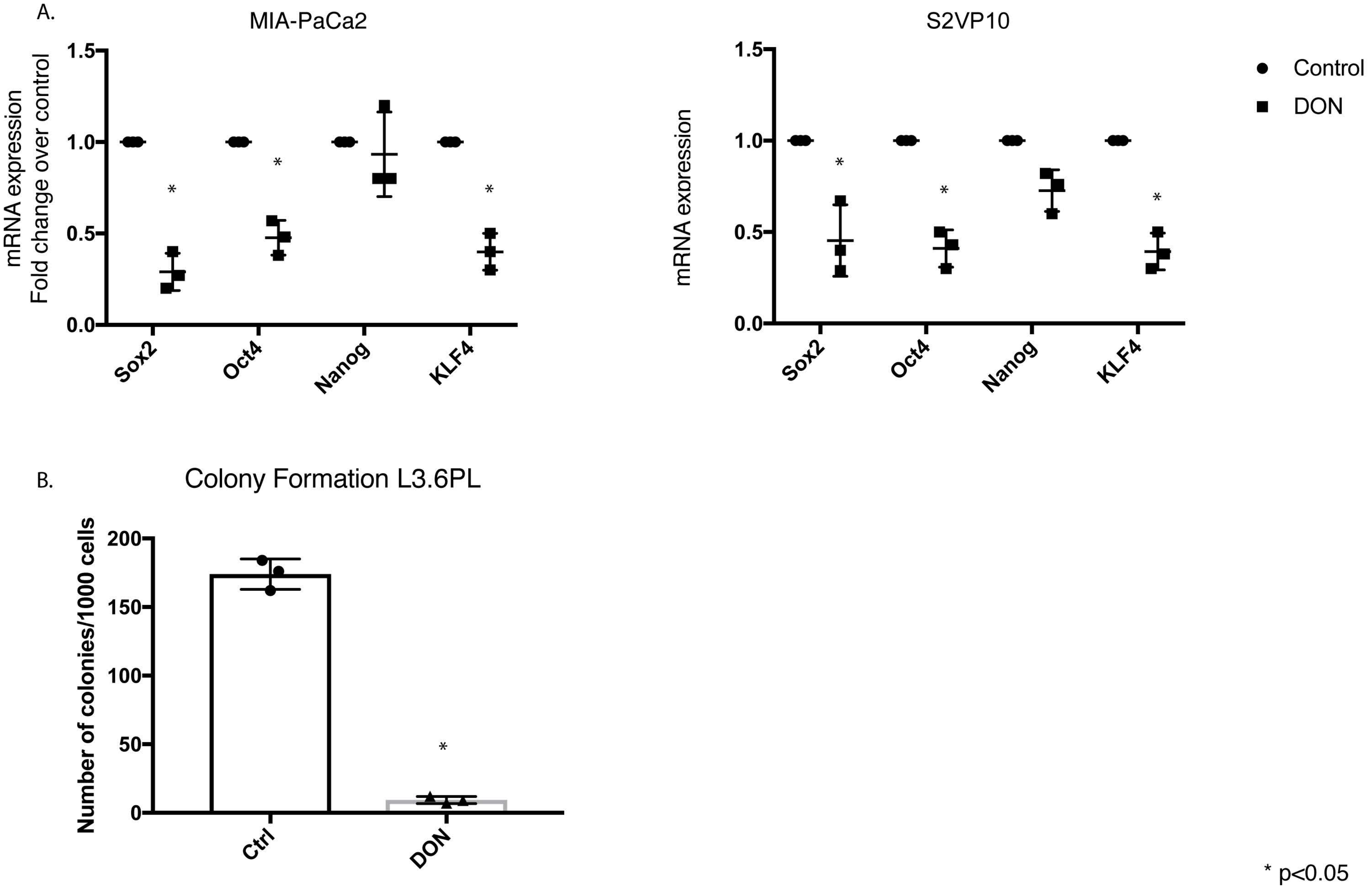


Suplementary Figure 1

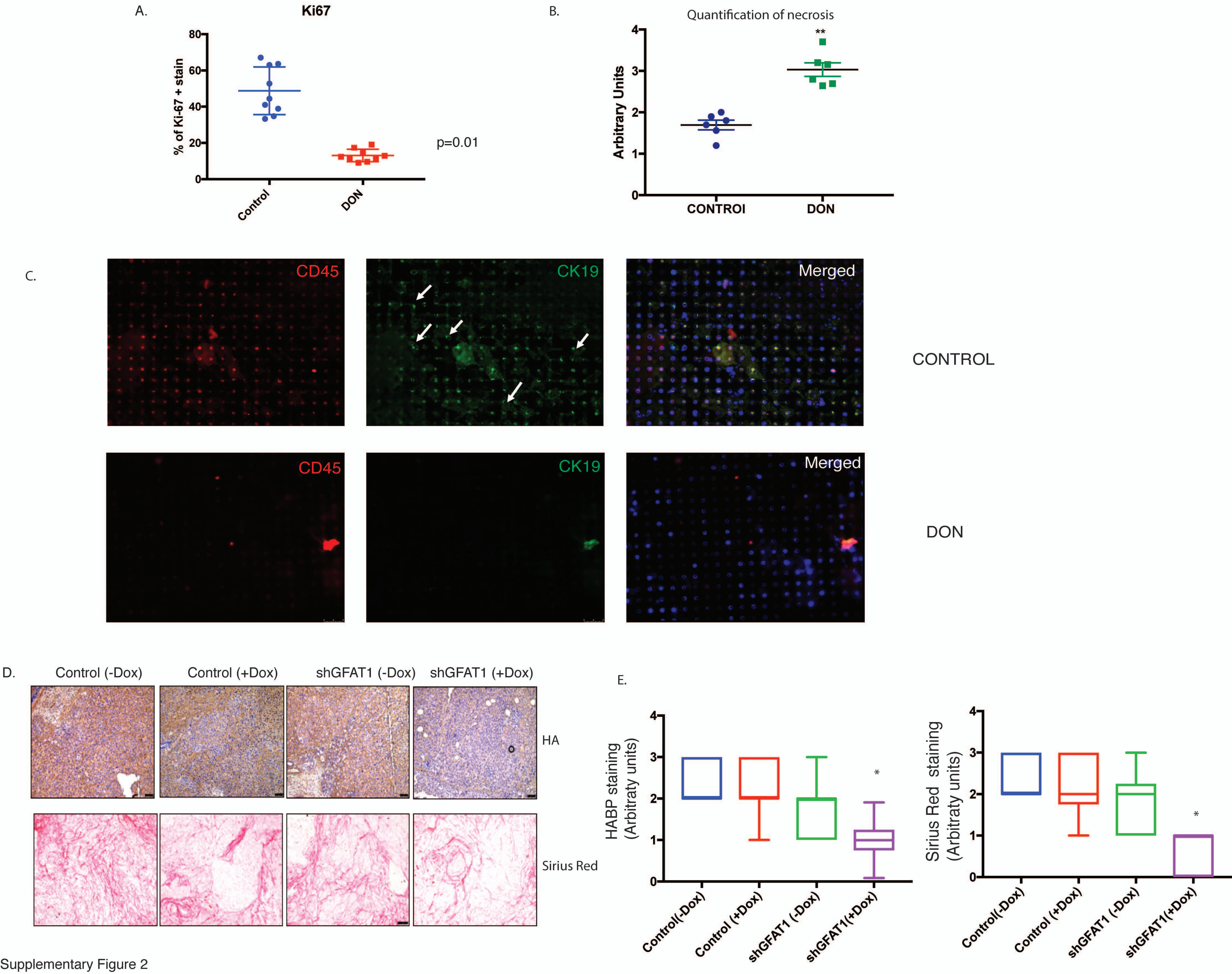


Supplementary Figure 2

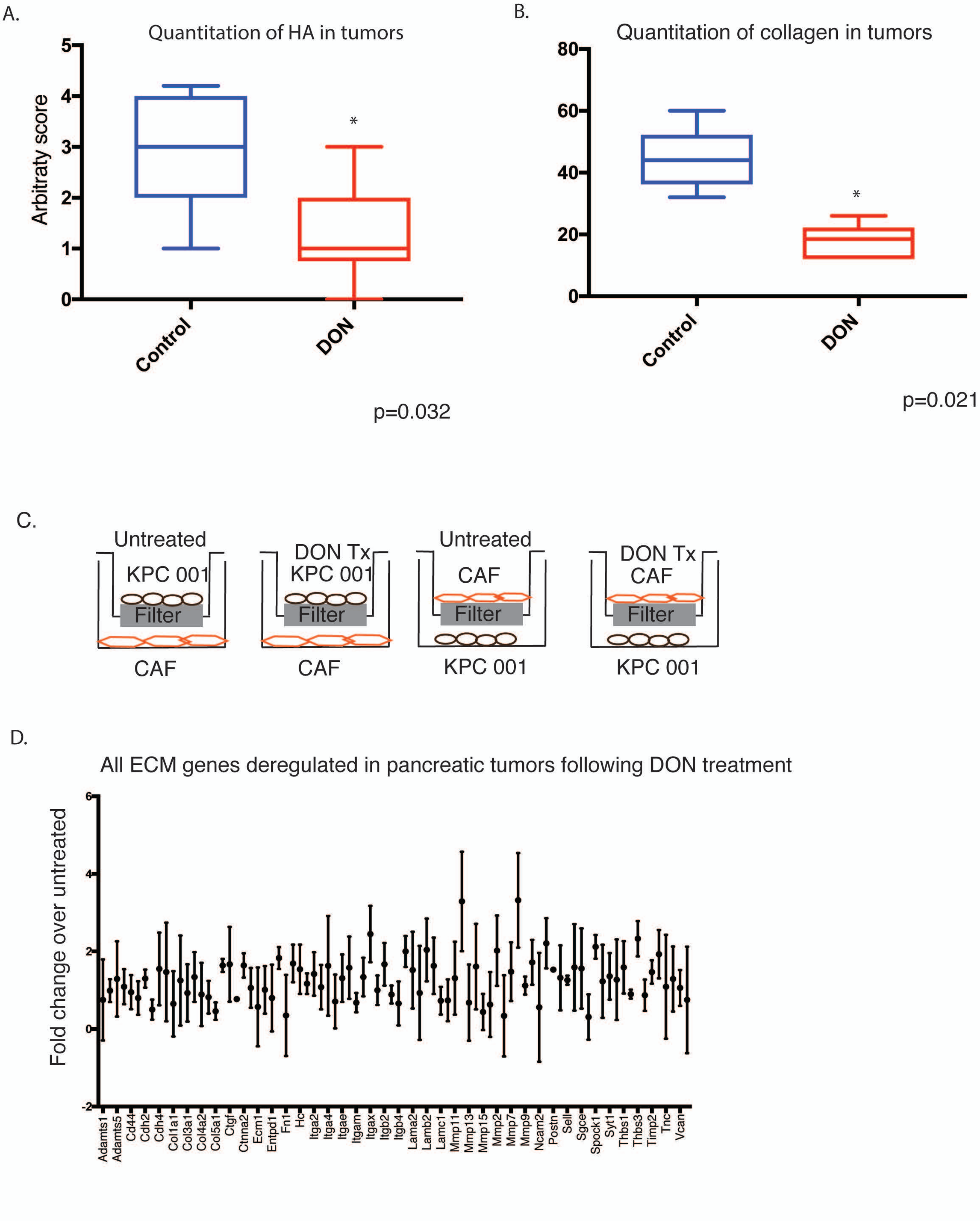


SUPPLEMENTARY FIGURE 3

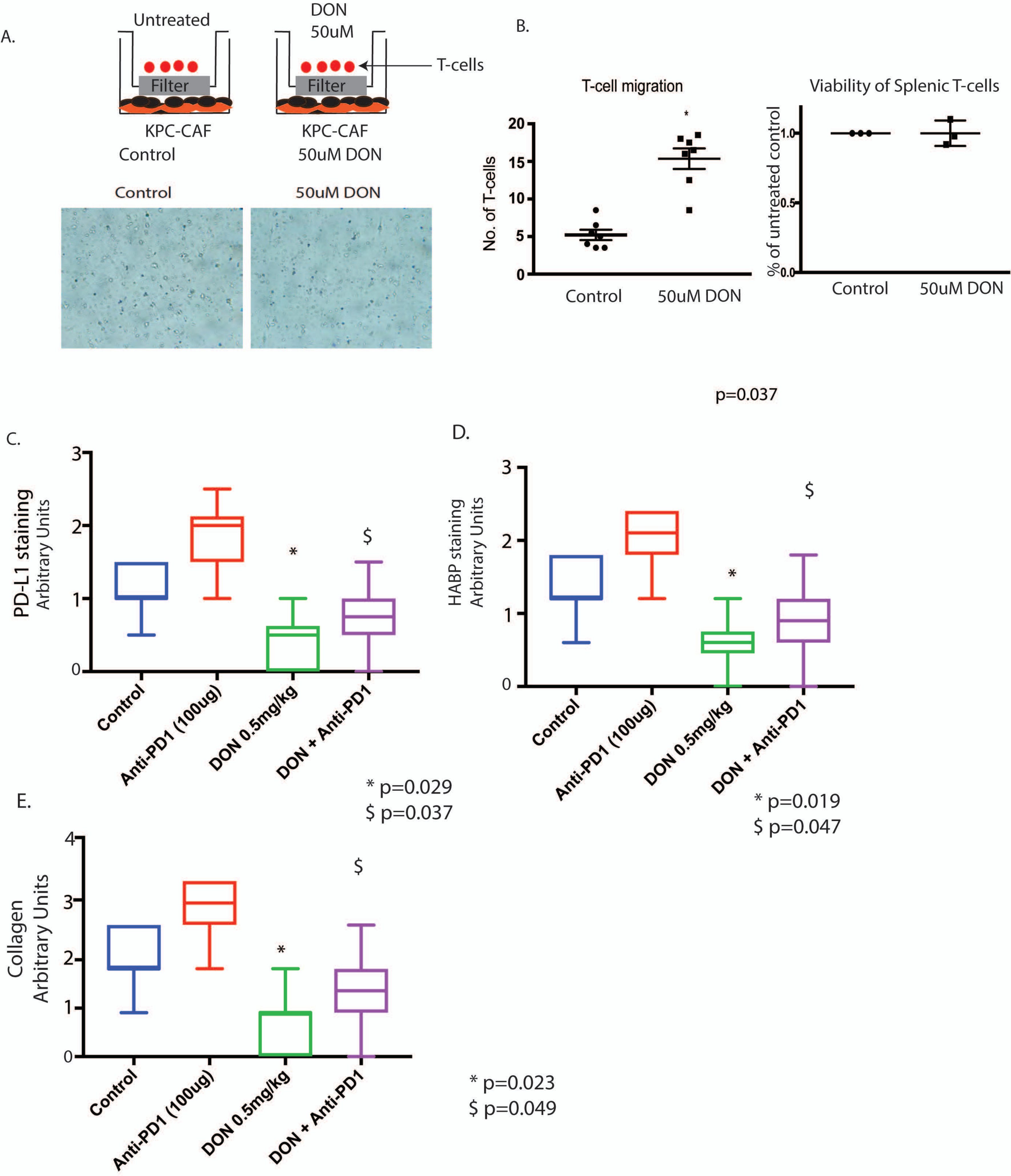
A.
Untreated
DON
50uM
T-cells
Filter
Filter
KPC-CAF
Control
KPC-CAF
50uM DON
Control
50uM DON
B.
T-cell migration
No. of T-cells
20
15
10
5
0
*
Control
50uM DON
Viability of Splenic T-cells
% of untreated control
1.0
0.5
0.0
Control
50uM DON
p=0.037
C.
PD-L1 staining
Arbitrary Units
3
2
1
0
*
$
Control
Anti-PD1 (100ug)
DON 0.5mg/kg
DON + Anti-PD1
* p=0.029
$ p=0.037
D.
HABP staining
Arbitrary Units
3
2
1
0
*
$
Control
Anti-PD1 (100ug)
DON 0.5mg/kg
DON + Anti-PD1
* p=0.019
$ p=0.047
E.
Collagen
Arbitrary Units
4
3
2
1
0
*
$
Control
Anti-PD1 (100ug)
DON 0.5mg/kg
DON + Anti-PD1
* p=0.023
$ p=0.049

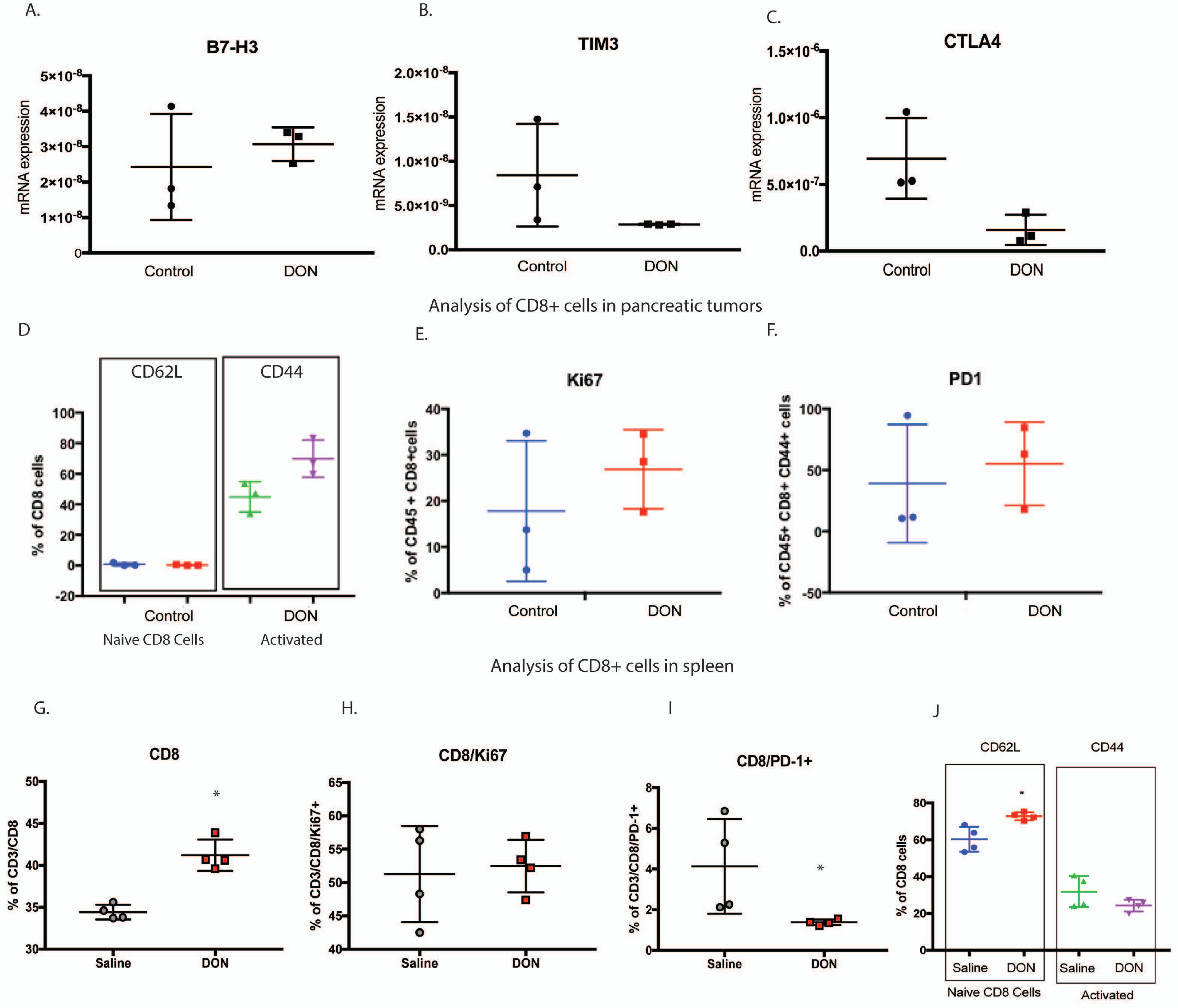
A.
B7-H3
mRNA expression
Control
DON
B.
TIM3
mRNA expression
Control
DON
C.
CTLA4
mRNA expression
Control
DON
Analysis of CD8+ cells in pancreatic tumors
D
CD62L
CD44
% of CD8 cells
Control
DON
Naive CD8 Cells
Activated
E.
Ki67
% of CD45 + CD8+cells
Control
DON
F.
PD1
% of CD45+ CD8+ CD44+ cells
Control
DON
Analysis of CD8+ cells in spleen
G.
CD8
% of CD3/CD8
Saline
DON
H.
CD8/Ki67
% of CD3/CD8/Ki67+
Saline
DON
I
CD8/PD-1+
% of CD3/CD8/PD-1+
Saline
DON
J
CD62L
CD44
% of CD8 cells
Saline
DON
Saline
DON
Naive CD8 Cells
Activated
